# Natural language processing captures memory content associated with shared neural patterns at encoding

**DOI:** 10.1101/2025.09.02.672707

**Authors:** June-Kyo Kim, Charan Ranganath, Alexander Barnett

## Abstract

People can experience the same event yet form distinct memories shaped by individual interpretations. Prior research shows that multivariate activity patterns in the Default Mode Network (DMN) are correlated across individuals during shared experiences, suggesting a role in representing high-level event features. However, it remains unclear whether these shared neural patterns reflect similarity in subsequent memory content. Here, we examined whether memory similarity correlates with intersubject spatial patterns in the DMN. Using topic modeling, we transformed verbal recall into vectors of latent topics to quantify memory similarity across participants. Twenty-four individuals watched and recounted two cartoon movies during fMRI scanning. We found that greater similarity in recalled content was associated with stronger shared activation patterns at encoding, particularly in the posterior medial cortex. These findings highlight the utility of natural language processing tools in linking memory representations to brain activity and underscore the DMN’s role in encoding and interpreting complex event features.

---

Narratives unify shared experiences between people and are thought to be adaptive for group cooperation (Smith et al., 2017). However, even when watching or reading the same narrative passage, people may notice unique details or find special significance in some elements of the story compared to others. These shared and unique elements shape how we remember our experiences.

Previous studies have found that encoding and retrieval of narratives drives shared multivariate patterns in default mode network (DMN) regions among people watching or recalling the same stories (Bird et al., 2015; J. Chen et al., 2017). These patterns are event specific suggesting that memory representations for real-world events are spatially organized in a manner that is generalizable across brains (Yeshurun et al., 2021).

Neocortical DMN regions are thought to represent higher-order, abstracted event models of ongoing situations (Fernandino & Binder, 2024; Franklin et al., 2020; Reagh & Ranganath, 2018). Further, this collection of regions has been shown to functionally couple with each other and the hippocampus at rest to form a large-scale intrinsic network (Barnett et al., 2021; Buckner et al., 2008) and interactions of these regions with the hippocampus during encoding and retrieval are thought to reflect the storage and reactivation of event-specific information (Barnett et al., 2024; Franklin et al., 2020; Lu et al., 2022; Mishra et al., 2025).

Prior research has shown that individuals who share the same interpretation of a narrative had greater inter-subject synchrony of their timecourses in the majority of the DMN (Nguyen et al., 2019; Yeshurun et al., 2017), and have greater correlations in their multivariate patterns (Zadbood et al., 2022) compared to those with an opposing interpretation. This work has linked the medial prefrontal cortex (mPFC), posterior medial cortex (PMC) and anterior temporal lobes (ATL) with higher order event representation as multivariate neural patterns during recall were more correlated when participants recalled events with the same interpretation (Zadbood et al., 2022). These findings are in line with theories that the DMN represents abstract event models (Franklin et al., 2020; Reagh & Ranganath, 2018).

These previous studies compared participants in discrete groups by experimentally manipulating their interpretations of events (Yeshurun et al., 2017; Zadbood et al., 2022). These manipulations involved either selectively providing or withholding information to bias the meaning of events. However, such approaches risk oversimplifying the multifaceted nature of narrative memory. For example, one pair of subjects may share many aspects of their recall, another pair may share a few, and another pair may have opposing views of an event. Recent advances in natural language processing models have provided powerful tools for analyzing the language people use during recall (Nguyen et al., 2019). These models can uncover latent themes in spoken or written recollections, allowing researchers to identify both shared and unique aspects of remembered content (Heusser et al., 2021). By examining which combinations of latent themes are common across pairs of participants, this approach enables a more nuanced assessment of the similarity in their memory representations.

In this study, we used natural language processing and fMRI to examine how multivariate event patterns in the DMN are shared during encoding and recall, depending on the degree of similarity in which participants remembered events. Twenty-four participants watched and recalled two cartoon movies during fMRI scanning. To quantify recall similarity, we used topic modeling to estimate the latent topics in each participant’s recall and measured the similarity of these topic patterns between participants. We calculated intersubject pattern similarity (ISPS) in regions of the DMN and examined whether ISPS at encoding and recall was associated with recall similarity. We hypothesized that people who recall events more similarly to each other would show stronger ISPS in DMN regions, particularly the mPFC and PMC – regions that have been heavily implicated in representing higher order abstractions of events (Audrain & McAndrews, 2022; Baldassano et al., 2018; Reagh & Ranganath, 2018; Zadbood et al., 2022). We also examined how recall similarity related to simpler scoring measures of memory content such as the total number of details recalled and the number of intrusions. We observed that ISPS at encoding was strongly correlated with recall similarity, meaning that people who had shared multivariate patterns in the DMN as they viewed events tended to recall similar content.

## Methods

### Dataset

Our dataset consisted of 24 healthy, right-handed young adults with no history of neurological issues who watched two 16.5 minute custom-made cartoon movies and then freely recalled them in the scanner (Barnett et al., 2024).

### MRI acquisition

MRI data was collected at both encoding and recall using a 3T Siemens Skyra scanner system with a 32-channel head coil. Two T1-weighted structural images were acquired using a magnetization prepared rapid acquisition gradient-echo (MPRAGE) pulse sequence (TR = 1900 ms) and functional images were acquired using a gradient EPI sequence (TR = 1220 ms). For detailed scanning parameters see Barnett et al. (2024). Functional images were collected during the viewing of two 16.5-minute animated movies and during verbal free recall. During viewing, participants were instructed to watch the movies as if they were a television show they were interested in. Without cues, participants freely recalled the movies aloud using an MRI compatible microphone. They were instructed to “In as much detail as possible tell me everything you can remember about the last movie we showed you. Try to recount the events in the original order in which they occurred. Completeness and detail are more important than temporal order. If at any point you realize that you have missed something, return to it. Try to describe EVERY detail that you have about the movie you just watched, even if it seems irrelevant”.

### Recall scoring

Audio recall for each participant was transcribed and scored. Details were defined as “a unique occurrence, observation, or thought, typically expressed as a grammatical clause. Additional information in the clause was scored separately” as defined by Levine et al. (2002). Only details that were verifiably true according to the events depicted in the movies were considered. Recall transcripts were then inspected and labeled by blocks of time corresponding to recall of a given event allowing us to tabulate the number of verifiable details recalled for each event. In some cases, participants would recall a detail from another event during the recounting of an event. These details were classified as intrusions. For additional information on scoring and labeling procedures, see ref Barnett et al., 2024.

### Preprocessing

Initial preprocessing was done using fMRIPrep (Esteban et al., 2019). See Barnett et al., 2024 for full details. Anatomical preprocessing included skull stripping, brain tissue segmentation, cortical surface reconstruction using FreeSurfer (Fischl, 2012), and volume-based spatial normalization to standard space. Functional images were corrected for susceptibility distortion and were co-registered to the T1w reference image and then resampled into standard space. Regressors of no interest including six-degree motion parameters and motion outliers (framewise displacement > 0.5 mm, DVARS > 3) were regressed out of the data and the data was high-pass filtered to remove frequencies below.001Hz all in one step using tproject (Cox & Hyde, 1997) in nipype (Gorgolewski et al., 2011).

### Intersubject pattern similarity analysis

To examine multivoxel patterns, we constructed 10 ROIs by combining spatially contiguous regions of the DMN that were previously found to cluster into DMN subnetworks using intrinsic functional connectivity (Barnett et al., 2021) from the HCP-MMP1.0 atlas (Glasser et al., 2016; Horn, 2016). The 10 ROIs include the PMC, medial temporal lobe (MTL), precuneus (PreC), ventral mPFC (vmPFC), dorsal mPFC (dmPFC), ATL, ventral lateral parietal (vLP), dorsal lateral parietal (dLP), anterior lateral parietal (aLP), and a control region, the somatomotor cortex (SM) that we did not predict would track abstract internal event representations (**Figure 3A**). The combination of regions for each ROI can be found in Supplemental Table 1. The corrected voxel timecourses of each region were extracted then z-scored across the entire run (**Figure 1**). Mean voxel event activity was estimated by modeling the data with time blocks for each event convolved with AFNI’s hemodynamic response function (Cox & Hyde, 1997). A multivoxel activity pattern was created per event, per subject, and per region using linear regression to estimate the mean event activity. For every pair of participants, we calculated the Pearson’s correlation between event activity patterns of each matching event for every region of interest. These correlation values were Fisher z-transformed and used as the dependent variable, ISPS, in linear mixed models described below.

**Figure 1.**
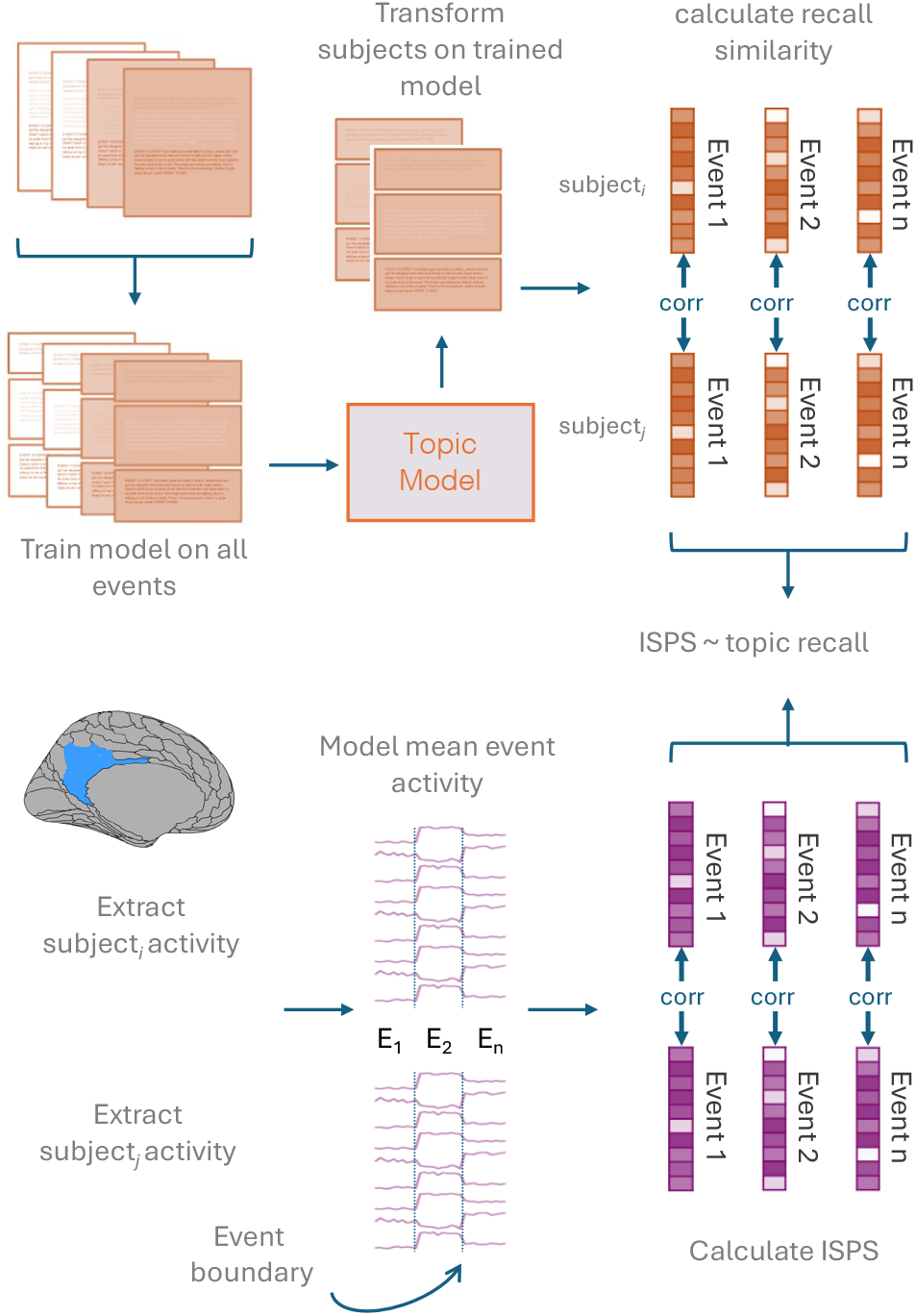
Overview of analysis pipeline. TOP: The preprocessed recall transcripts for each participant were divided into separate documents for each event. The collection of documents was then used to train a topic model to identify latent topics. Each event from each participant was then transformed into a vector of topic proportions that reveal the combination of topics present in the recalled event using the trained topic model. To calculate topic similarity, we correlated the vector of topic proportions from matching events between participants and Fisher z-transformed the correlation. BOTTOM: To examine intersubject pattern similarity (ISPS), multivoxel activity was extracted from each ROI and was modeled to find the mean multivoxel pattern for each event and each participant. We then correlated these multivoxel patterns for matching events between pairs of participants and Fisher-z transformed the correlation to get ISPS.

### Natural language processing

To quantify the similarity of people’s memory recall, the participant recall transcripts (corpuses) were used to perform topic modeling. We performed initial preprocessing including lemmatization and removing stop words using scikit-learn 1.4.1 (Pedregosa et al., 2011), manually removed punctuation, and standardized capitalization as done in Heusser et al. (2021). Additionally, we normalized the spelling of names or places in transcripts. For example, one participant’s recall may have been transcribed to call the main character Arlene and another participant’s recall may have been transcribed as Ailene. In this situation, we would convert all mistranslated names to the standard character’s name, Arlene. For each participant, we divided the recall transcript into separate based on the event that was being described. Independent raters read through each transcript and delineated the moments when a participant was referencing a specific event as defined in our previous paper (Barnett et al., 2024). Each delineated collection of recall for events served as “documents” in topic modeling.

Using methods developed by Heusser et al. (2021), we trained our model using these recall documents with scikit-learn 1.4.1, called from HyperTools (Pedregosa et al., 2011). Broadly, we used the CountVectorizer class, to transform the words from the collection of documents into a vector of word counts yielding a document-by-word count matrix. For the collection of documents, we submitted the preprocessed recall transcripts for each event and each participant (Figure 1). We used LatentDirichletAllocation class to train the topic model to the word count matrices, which identified latent themes within the documents. We could then apply the model to the data of each participant to evaluate the presence of each latent theme within each event, resulting in an event-by-topic proportion matrix per participant. Each event vector in these matrices describes the mixture of discovered topics (latent themes) present within the event. Using the event-by-topic proportion matrix as a proxy for recalled content, the similarity of recall per event between participants was estimated by correlating the event topic vectors between every pair of participants in the group in a pairwise fashion for matching events. All correlations were Fisher z-transformed to make a measurement we call *topic similarity*.

When training a topic model, certain parameters can affect the model, like the number of topics (*K*). Given that we didn’t have a strong prior regarding the number of topics, we performed a parameter sweep from 5 - 55 topics. Similarly, the fitting of the topic model will vary based on a random seed, resulting in slightly different models each time. To ensure that our findings are not sporadic, we also fit the model 100 times using a new random seed for each *K* number of topics and averaged the similarity to bootstrap our findings and used the average model output. The ‘batch’ learning method was used, and all other parameters were kept as default. The resulting topic similarity values were averaged across the range of topic numbers.

### Statistical Analyses

First, we examined how the topic similarity between participants for an event related to the number of details recalled from that event and the number of intruding details (intrusions) that were present in the recall of that event. We fit a linear mixed model to predict topic similarity from the number of details recalled using a fully crossed random effects structure (G. Chen et al., 2017) using lmer (Bates et al., 2015) in R 4.4.1 (R Core Team, 2021). In this structure, each topic similarity value in the model comes from a pair of participants *i* and *j*, where *i* ≠ *j*. Random effects are modeled for both the first and second participants in the pairing. Since the random effects are fully crossed, each participant appears in both the first random effect and the second random effect. However, this also leads to a data doubling in the model which is later corrected by discounting the degrees of freedom in the standard error and the degrees of freedom for the t-test and p-value calculations in line with (G. Chen et al., 2017) by dividing the number of observations by 2 and subtracting the number of fixed effects. Since topic similarity is calculated by a pair of participants, participant *i* and participant *j*, a linear mixed model was fit to predict topic similarity from the details recalled from participant *i* and the details recalled from participant *j* in line with (G. Chen et al., 2017). A separate linear mixed model was fit to predict topic similarity from intrusions in the same manner.

Next, a separate set of linear mixed models were fit to predict ISPS from topic similarity for each ROI at both encoding and retrieval with a fully crossed random effect approach. To determine whether ISPS was driven more by topic similarity or recalled details, we fit a separate set of models that included both ISPS and recalled details (for both participants) in the model to predict ISPS. When running the fully crossed linear mixed models for the recall ISPS we used the L-BFGS-B optimizer from optimx (Nash & Varadhan, 2011) to allow for more flexible model convergence. Also, when predicting recall ISPS, we included the number of TRs that elapsed as a participant described an event as a regressor of no interest to ensure that relationships between topic similarity and ISPS are not simply due to having more data to estimate multivoxel patterns. Significance was assessed at p <.05, FDR corrected and all predictors were mean centered.

## Results

### Similarity of topic patterns and scored details

We tested if recall similarity measured by topic similarity captured similar content as human raters to examine whether topic modeling accurately captured memory performance in terms of the number of details recalled. It is not strictly necessary that a greater number of recalled details should relate to higher topic similarity, but if participants have a tendency to recall similar latent themes, then recall of more information might allow for more precise estimation of topic proportions and higher similarity. Conversely, participants who mix in details from other events (intrusions) should have lower similarity to the rest of the group (unless everyone showed a tendency to have specific intrusions).

A linear mixed model revealed that event topic similarity predicted the number of event details that were recalled, (t(3994) = 11, p <.0001), and conversely, event topic similarity was negatively predicted by intrusions (t(3994) = -4.35, p <.0001), as expected. This demonstrates that the topics produced from the topic models are sensitive to the content of retrieval in an event-specific manner. When participants recall more true details and fewer intrusions, they show greater similarity (**Figure 2**).

**Figure 2.**
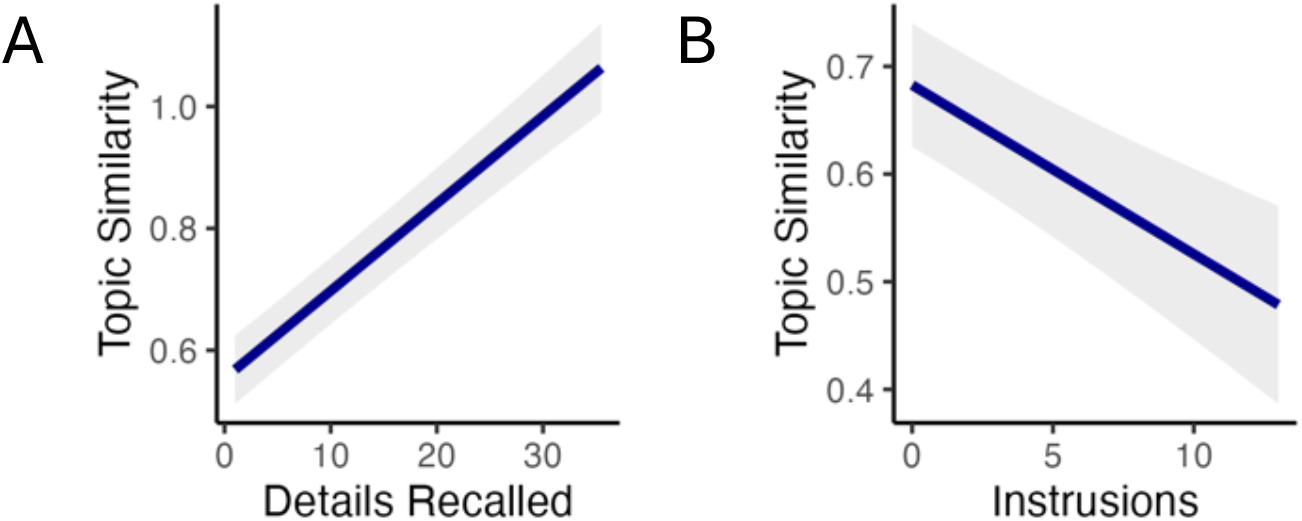
**A**. Topic similarity between participants increases with more details recalled. Line plots depict the relationship between topic similarity and details recalled. The shaded area around the line represents the standard error of the estimate. **B**. Greater number of intrusions related to decreased topic similarity. Line plots depict the relationship between topic similarity and the number of intrusions. The shaded area around the line represents the standard error of the estimate.

### Relationship between topic similarity and intersubject pattern similarity

Having observed that topic similarity is related to manual human scoring, we then asked whether topic similarity was associated with neural ISPS using linear mixed models. Given that enhanced intersubject correlations (temporally correlated timecourses between participants) during encoding has been associated with better memory performance (Halpern et al., 2023; Hasson et al., 2008; Koch et al., 2020), we also included the number of details recalled in the same model to determine whether content similarity related to ISPS independent of just having better memory. We observed that topic similarity was significantly positively associated with encoding ISPS in nearly all DMN cortical regions, but not the control ROI (**Figure 3B**) (PMC: t(3993)=6.92, p<.0001; dmPFC: t(3993) = 4.5; p <.0001; ATL: t(3993) = 4.4, p <.0001; vmPFC: t(3993) = 3.02, p =.005; MTL: t(3993) = 2.63, p =.011; PreC: t(3993) = 3.81, p =.0004; dLP: t(3993) = 2.73, p =.009; aLP: t(3993) = 3.75, p =.0004). These results show that participants who had similar multivoxel patterns at encoding, tended to recall a similar collection of topics at retrieval. Test statistics for all regions are presented in Supplemental Table 2.

**Figure 3.**
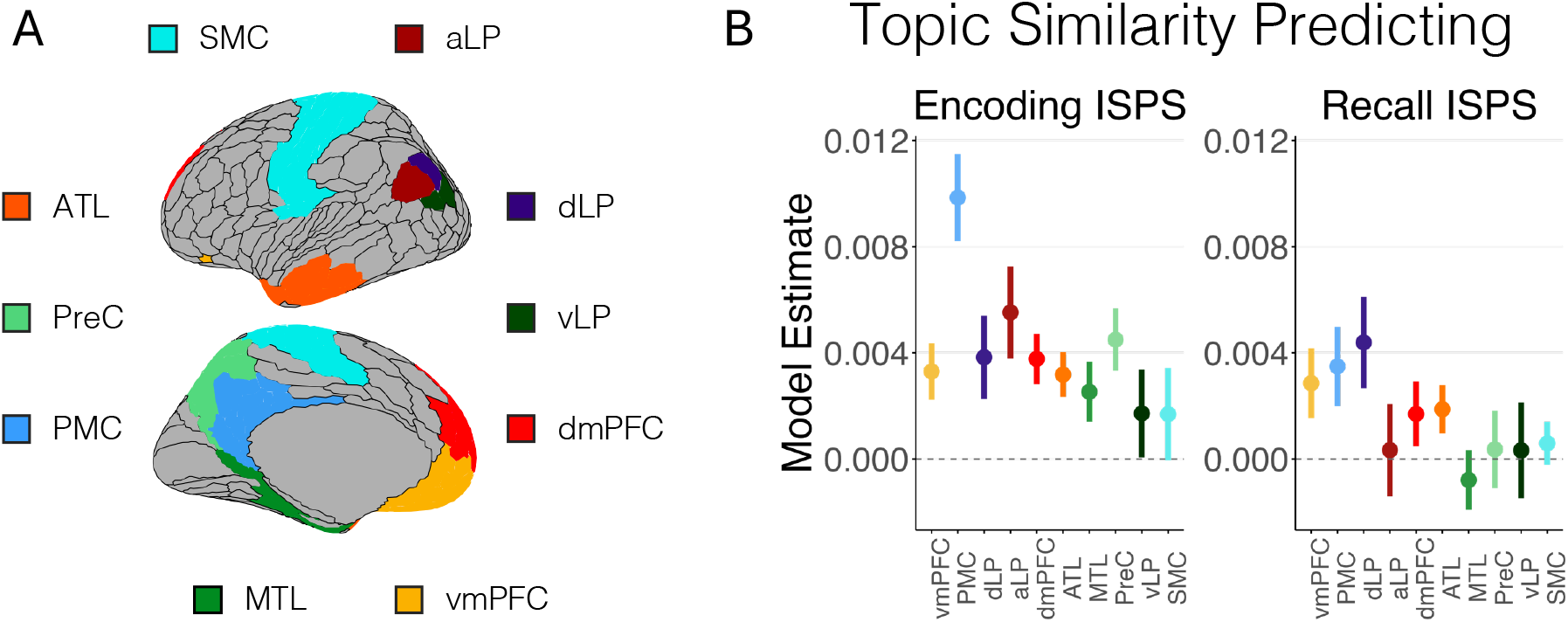
Topic similarity relates to ISPS. **A**. Depiction of ROIs on inflated brain **B**. Point plot showing the estimated model slopes for topic similarity when predicting intersubject pattern similarity (ISPS) at encoding (left) and recall (right). Error bars represent the standard error of the estimate. ATL, anterior temporal lobe; aLP, anterior lateral parietal cortex; dLP, dorsal lateral parietal cortex; dmPFC, dorsomedial prefrontal cortex; MTL, medial temporal lobe; PMC, posterior medial cortex; PreC, precuneus; SMC, somatomotor; vLP, ventral lateral parietal; vmPFC, ventromedial prefrontal cortex.

On the other hand, when looking at recall activity, topic similarity was positively associated with recall intersubject patterns only at uncorrected levels in the PMC, ATL, vmPFC, and dLP (**Figure 3B**) (PMC: t(3993)= 2.49, p =.01 uncorr; ATL: t(3993)= 2.37, p =.02 uncorr; vmPFC: t(3993)= 2.21, p =.03 uncorr; dLP: t(3993)= 2.53, p =.01 uncorr). Test statistics for all regions are presented in Supplemental Table 3.

## Discussion

When experiencing the same event, people can often remember it in different ways. Here, we sought to examine what areas of the brain relate to the combination of features we recall. Using natural language processing tools, we quantified the similarity of memory content for naturalistic events between individuals. We found that the similarity of retrieved content tracked with the level of detail for those memories and was inversely related to the number of intrusions. Further, recall similarity was reliably associated with shared neural patterns across participants— even after accounting for the amount of detail recalled. Specifically, shared neural patterns at encoding were associated with similar memory content during recall and this was particularly true in posterior medial, lateral parietal, medial prefrontal and medial temporal cortices. This association provides evidence that the neural patterns in these regions during the initial encoding of an event influence the subsequent content recalled during retrieval.

Natural language processing and recently developed large language models are increasingly being explored to characterize memory retrieval and event cognition (Fenerci et al., 2025; Georgiou et al., 2025; Heusser et al., 2021; Lee & Chen, 2022; Michelmann et al., 2025; Panela et al., 2025; Su et al., 2024). This work has shown that the abstract features generated from topic modeling or text embeddings can be used to segment narratives into discrete events similar to the way humans segment events (Heusser et al., 2021; Michelmann et al., 2025; Panela et al., 2025) and have linked changes in semantic content with neural state changes in DMN regions (Yang et al., 2023). Further, these models are beginning to be used to score recall accuracy showing concordance with human raters (Mistica et al., 2024; Van Genugten & Schacter, 2024) as well as to examine the diversity of information retrieved (Sheldon et al., 2024). We trained a topic model on the recall transcripts generated by participants in the study to calculate a set of latent topics at the group level. We observed that when participants were able to generate more details, they also tended to show greater similarity in the pattern of topics used to describe a given event. However, and importantly, when participants generated intrusions (details from another event), we observed lower similarity in the pattern of topics used to describe the event suggesting that the topics discovered track the within event features.

Previous work has demonstrated that multivariate patterns within cortical regions of the DMN show event-specific patterns that shift at event boundaries during encoding (Baldassano et al., 2017; Wilford et al., 2025). These same patterns are reinstated at retrieval (J. Chen et al., 2017; Oedekoven et al., 2017), and the magnitude of reinstatement is proportional to the amount of hippocampal boundary activity observed when these patterns switch (Baldassano et al., 2017). These DMN regions are thought to represent the ongoing event model of the current situation (Franklin et al., 2020). That is to say, these regions are thought to integrate the current ongoing experience with prior event knowledge to form an abstracted representation that allows for interpretation and predictions of incoming information (Zacks, 2020). Indeed, when events share abstracted event information, multivariate patterns in these areas are more likely to generalize (Baldassano et al., 2018; Karagoz et al., 2023; Reagh & Ranganath, 2023). During encoding, interaction between the hippocampus and DMN at event boundaries seem to allow for the long term storage of these abstracted representations allowing for successful subsequent retrieval (Barnett et al., 2024; Mishra et al., 2025; Park et al., 2025). This degree of abstraction fits with topographic network analysis of the brain that has shown the DMN sits at the apex of a cortical hierarchy (Margulies et al., 2016) receiving abstracted external information from intermediary networks (Barnett et al., 2021). Further, as people attend and encode to incoming information, changes in semantic content coincide well with multivariate changes in DMN regions (Yang et al., 2023). Our findings, showing a relationship between abstracted topics and multivariate patterns in the DMN fit well with the notion that the DMN is capturing abstracted, higher-order information in service of event cognition. Collectively, this evidence points to the DMN’s role in combining incoming information with prior semantic knowledge to represent the ongoing interpretation of events, ultimately impacting the content recalled at retrieval.

Remarkably, multivariate patterns within the DMN at encoding and recall are shared between people (J. Chen et al., 2017; Oedekoven et al., 2017). Our work extends this to show that the degree of overlap in these patterns is associated with the content of subsequent retrieval. Previous work has shown that when two people watch the same stimulus, but have different interpretations, they will have more distinct patterns and dynamics in these regions compared to when they have the same representation (Sava-Segal et al., 2023; Yeshurun et al., 2017; Zadbood et al., 2022). Our findings are strongly concordant with these previous observations and capitalize on the memory representations generated by the participants to identify subject to subject and event to event variations in recall similarity. Our results extend on this previous work showing that the spatial patterns established during viewing in DMN regions, which relate to ongoing interpretation of events, are indicative of the subsequent memory representations that will be formed and retrieved. These effects are even more impressive given that they are found even after accounting for the detailedness of memory representations.

We did not observe significant relationships between recall similarity and recall ISPS— though they were in the expected direction based on prior work (Zadbood et al., 2022). One challenge facing recall ISPS is that the amount of fMRI data available from participants is often less than the amount of data at encoding (J. Chen et al., 2017). Additionally, participants often had higher motion during recall which could affect estimated patterns even after preprocessing. Recall also may contain additional processes, difficult to capture such as construction and elaboration (Conway, 2005) which engage the DMN and hippocampus in differing manners (Audrain et al., 2022; McCormick et al., 2015). Indeed, participants in our study would occasionally pause during recall, possibly indicating moments of memory search. While we used the presence of speech to determine timepoints for recalling specific events which likely occurs during elaboration, it is possible that other processes could contaminate or complicate the multivariate event patterns.

One challenge with natural language processing models is the multiple model parameters that can be explored. Additionally, the training of topic models can result in slightly different weightings and groupings each time due to the stochastic nature of the word by document decomposition. To account for these challenges, we averaged across a range of model parameters and model training iterations, as we had no strong hypotheses on the number of latent themes that would best model the recall content. Additionally, in our study, we relied on verbal free recall to estimate memory representation similarity. Given that people tend to have different levels of verbosity, we may be able to better represent latent themes present in some participants’ memory representations compared to others who speak more reservedly. This limitation is typical in free recall of naturalistic or autobiographical event studies, even when experimenter probing is provided to request more information (Lockrow et al., 2023). Future studies could ask specific probing questions about memories and interpretations surrounding aspects of events that may be subserved by different regions or subnetworks of the DMN (Andrews-Hanna et al., 2014; Barnett et al., 2021; Braga et al., 2019; DiNicola et al., 2020). For example, one could specifically ask about emotional tones or themes, the meaning of social interactions or sarcasm, or perceptual details. These focused questions could provide deeper insight into where individuals converge or diverge in their recollections and interpretations. This approach could also address a key limitation of topic modeling: the difficulty in interpreting the resulting latent topics. These topics of lack a clear semantic categorization or label. Asking targeted questions in future experiments could help link neural representations with specific types of content.

In conclusion, the present study demonstrates that activity patterns within the DMN during encoding predict the content people subsequently retrieve. These findings reinforce the role of the DMN in maintaining event model representations that reflect our ongoing understanding of an event and our eventual memory representations for that event. These findings also highlight the utility of topic modeling in providing insight in characterizing memory representations.

## Acknowledgements

The authors would like to thank Sam Audrain for helpful comments on the manuscript. JKK was supported by a Natural Sciences and Engineering Research Council (NSERC) Undergraduate Summer Research Award. This research was supported by an NSERC Discovery Grant (RGPIN-2023-05010) and a Fonds de Recherche du Quebec Chercheur Boursier (AJB) and by a Multi-University Research Initiative grant N00014-17-1-2961 from the US Office of Naval Research/Department of Defense (CR)

---

**Supplemental Table 1.** Topic similarity and Encoding ISPS.

| Glasser ROI name | hemisphere | ROI Combo |
| --- | --- | --- |
| RSC | L | PMC |
| 7m | L | PMC |
| POS1 | L | PMC |
| v23ab | L | PMC |
| d23ab | L | PMC |
| 31pv | L | PMC |
| 31pd | L | PMC |
| 31a | L | PMC |
| RSC | R | PMC |
| 7m | R | PMC |
| 23d | R | PMC |
| v23ab | R | PMC |
| d23ab | R | PMC |
| 31pv | R | PMC |
| 31pd | R | PMC |
| 31a | R | PMC |
| d32 | L | dmPFC |
| 9m | L | dmPFC |
| 8BL | L | dmPFC |
| 9p | L | dmPFC |
| 10d | L | dmPFC |
| 9a | L | dmPFC |
| d32 | R | dmPFC |
| 9m | R | dmPFC |
| 8BL | R | dmPFC |
| 9p | R | dmPFC |
| 10d | R | dmPFC |
| 9a | R | dmPFC |
| TGd | L | Lat temp |
| TE1a | L | Lat temp |
| TE2a | L | Lat temp |
| STSva | L | Lat temp |
| TE1m | L | Lat temp |
| TGd | R | Lat temp |
| TE1a | R | Lat temp |
| TE2a | R | Lat temp |
| STSva | R | Lat temp |
| TE1m | R | Lat temp |
| a24 | L | vmPFC |
| p32 | L | vmPFC |
| 10r | L | vmPFC |
| 47m | L | vmPFC |
| 10v | L | vmPFC |
| 10pp | L | vmPFC |
| OFC | L | vmPFC |
| 25 | L | vmPFC |
| s32 | L | vmPFC |
| pOFC | L | vmPFC |
| a24 | R | vmPFC |
| p32 | R | vmPFC |
| 10r | R | vmPFC |
| 47m | R | vmPFC |
| 10v | R | vmPFC |
| 10pp | R | vmPFC |
| OFC | R | vmPFC |
| 25 | R | vmPFC |
| s32 | R | vmPFC |
| pOFC | R | vmPFC |
| PreS | L | MTL |
| ProS | L | MTL |
| PeEc | L | MTL |
| PHA1 | L | MTL |
| PHA3 | L | MTL |
| PHA2 | L | MTL |
| PreS | R | MTL |
| ProS | R | MTL |
| PeEc | R | MTL |
| PHA1 | R | MTL |
| PHA3 | R | MTL |
| TF | R | MTL |
| PHA2 | R | MTL |
| PCV | L | POS_PreC |
| 7Pm | L | POS_PreC |
| 7Am | L | POS_PreC |
| 7PL | L | POS_PreC |
| DVT | L | POS_PreC |
| POS2 | L | POS_PreC |
| PCV | R | POS_PreC |
| 7Pm | R | POS_PreC |
| POS1 | R | POS_PreC |
| 7Am | R | POS_PreC |
| 7PL | R | POS_PreC |
| TPOJ3 | L | vLP |
| PGp | L | vLP |
| TPOJ3 | R | vLP |
| PGp | R | vLP |
| PGs | L | dLP |
| PGs | R | dLP |
| PGi | R | dLP |
| PFM | L | aLP |
| PGi | L | aLP |
| 4 | L | SMC |
| 3b | L | SMC |
| 5m | L | SMC |
| 5L | L | SMC |
| 24dd | L | SMC |
| 24dv | L | SMC |
| 1 | L | SMC |
| 2 | L | SMC |
| 3a | L | SMC |
| 6d | L | SMC |
| 6mp | L | SMC |
| 6v | L | SMC |
| OP4 | L | SMC |
| OP1 | L | SMC |
| OP2-3 | L | SMC |
| FOP2 | L | SMC |
| Ig | L | SMC |
| 4 | R | SMC |
| 3b | R | SMC |
| 5m | R | SMC |
| 5L | R | SMC |
| 24dd | R | SMC |
| 24dv | R | SMC |
| 1 | R | SMC |
| 2 | R | SMC |
| 3a | R | SMC |
| 6d | R | SMC |
| 6mp | R | SMC |
| 6v | R | SMC |
| OP4 | R | SMC |
| OP1 | R | SMC |
| OP2-3 | R | SMC |
| FOP2 | R | SMC |
| lg | R | SMC |

ATL, anterior temporal lobe; aLP, anterior lateral parietal cortex; dLP, dorsal lateral parietal cortex; dmPFC, dorsomedial prefrontal cortex; MTL, medial temporal lobe; PMC, posterior medial cortex; PreC, precuneus; SMC, somatomotor; vLP, ventral lateral parietal; vmPFC, ventromedial prefrontal cortex.

**Supplemental Table 2.** Topic similarity and Encoding ISPS. Relationship between Topic similarity and Encoding ISPS (accounting for details) DF = 3993

| ROI | t-value | p-value | FDR corrected |
| --- | --- | --- | --- |
| PMC | 6.92 | 6.7e-9 | **5.41e-11** |
| dmPFC | 4.5 | .000007 | **.00004** |
| ATL | 4.4 | .000016 | **.00004** |
| vmPFC | 3.02 | .0025 | **.005** |
| MTL | 2.63 | .0085 | **.01** |
| PreC | 3.81 | .00014 | **.0004** |
| vLP | 1.48 | .14 | .15 |
| dLP | 2.73 | .0063 | **0.009** |
| aLP | 3.75 | .00018 | **0.0004** |
| SMC | 1.45 | .15 | .15 |

ATL, anterior temporal lobe; aLP, anterior lateral parietal cortex; dLP, dorsal lateral parietal cortex; dmPFC, dorsomedial prefrontal cortex; MTL, medial temporal lobe; PMC, posterior medial cortex; PreC, precuneus; SMC, somatomotor; vLP, ventral lateral parietal; vmPFC, ventromedial prefrontal cortex.

**Supplemental Table 3.** Topic similarity and Recall ISPS. Relationship between Topic similarity and Recall ISPS (accounting for details) DF = 3993

| ROI | t-value | p-value | FDR corrected |
| --- | --- | --- | --- |
| PMC | 2.48 | .013 | 0.06 |
| dmPFC | 1.62 | .11 | 0.21 |
| ATL | 2.37 | .017 | 0.06 |
| vmPFC | 2.21 | .027 | 0.07 |
| MTL | -.47 | .64 | 0.71 |
| PreC | .68 | .5 | 0.66 |
| vLP | .22 | .82 | 0.82 |
| dLP | 2.53 | .012 | 0.06 |
| aLP | .63 | .53 | 0.66 |
| SMC | .98 | .33 | 0.55 |

ATL, anterior temporal lobe; aLP, anterior lateral parietal cortex; dLP, dorsal lateral parietal cortex; dmPFC, dorsomedial prefrontal cortex; MTL, medial temporal lobe; PMC, posterior medial cortex; PreC, precuneus; SMC, somatomotor; vLP, ventral lateral parietal; vmPFC, ventromedial prefrontal cortex.

